# SLC2A knockout mice deficient in ascorbic acid synthesis recapitulate aspects of arterial tortuosity syndrome and display mitochondrial respiration defects

**DOI:** 10.1101/862425

**Authors:** Annekatrien Boel, Joyce Burger, Marine Vanhomwegen, Aude Beyens, Marjolijn Renard, Sander Barnhoorn, Christophe Casteleyn, Dieter P. Reinhardt, Benedicte Descamps, Christian Vanhove, Ingrid van der Pluijm, Paul Coucke, Andy Willaert, Jeroen Essers, Bert Callewaert

## Abstract

Arterial tortuosity syndrome (ATS) is a recessively inherited connective tissue disorder, mainly characterized by tortuosity and aneurysm formation of the major arteries. ATS is caused by loss-of-function mutations in *SLC2A10*, encoding the facilitative glucose transporter GLUT10. Former studies implicate GLUT10 in transport of dehydroascorbic acid, the oxidized form of ascorbic acid (AA). Mouse models carrying homozygous *Slc2a10* missense mutations do not recapitulate the human phenotype. Since mice, in contrast to humans, are able to intracellularly synthesize AA, we generated a novel ATS mouse model, deficient for *Slc2a10* as well as *Gulo*, which encodes for L-gulonolactone oxidase, an enzyme catalyzing the final step in AA biosynthesis in rodents. *Gulo;Slc2a10* knock-out mice show mild phenotypic anomalies, which were absent in single knock-out controls. While *Gulo;Slc2a10* knock-out mice do not fully phenocopy human ATS, histological and immunocytochemical analysis revealed compromised extracellular matrix formation. TGFβ signaling remained unaltered, while mitochondrial function was compromised in smooth muscle cells derived from *Gulo;Slc2a10* knock-out mice. Altogether, our data add evidence that ATS is an ascorbate compartmentalization disorder, but additional factors underlying the observed phenotype in humans remain to be determined.

## Introduction

Arterial Tortuosity Syndrome (ATS, MIM#208050) is a rare autosomal recessive connective tissue disorder, characterized by severe patterning defects in blood vessel development. Main features of the disease include elongation, tortuosity, and stenoses of the large and middle-sized arteries, with an increased risk for aneurysm formation and ischemic events [1–3]. Patients may present with additional connective tissue-related features, such as *cutis laxa*, diaphragmatic hernias, joint laxity, distinctive craniofacial malformations and a marfanoid habitus. Finally, corneal thinning causing pellucid corneas and keratoconus were recently described [2, 4]. The disease has shown to be variable in terms of severity, ranging from early mortality to mild manifestations in adulthood [1, 2, 5, 6]. Arterial histopathology shows disorganization and fragmentation of the elastic fibers [1, 7–10]. Cultured fibroblasts of ATS patients show disorganization of the actin cytoskeleton and multiple extracellular matrix (ECM) components, including fibronectin, fibrillin, type 3 collagen, type 5 collagen and decorin [11, 12].

ATS is caused by loss-of-function mutations in the *SLC2A10* gene, encoding the GLUT10 protein [3]. GLUT10 belongs to the GLUT family of facilitative transporters, consisting of 14 transmembrane proteins, enabling the transport of monosaccharides, polyols, and other small carbon compounds across the membranes of eukaryotic cells [13]. GLUT members show specific functionalities, but some functional redundancy has been observed. Recent evidence identified GLUT10 as a transporter of dehydroascorbic acid (DHA), the oxidized form of ascorbic acid (AA), but its subcellular localization remains enigmatic and may change upon physiological states. GLUT10 has been shown to reside in mitochondria [14], the nuclear envelope [3], and/or the endoplasmic reticulum (ER) [15–17].

It remains to be solved how defective DHA transport results in a vasculopathy related to Loeys-Dietz [18] or Marfan Syndrome [19, 20], respectively caused by perturbed transforming growth factor beta (TGFβ) pathway signaling or altered extracellular matrix (ECM) proteins. In the absence of relevant vascular patient material, several ATS disease models have been developed. *Slc2a10* knockdown in a morpholino-based zebrafish model shows abnormal vascular patterning and mitochondrial respiration [21]. Two knock-in mouse models, carrying respectively a homozygous likely pathogenic G128E or S150F missense substitution in *Slc2a10* [22, 23], were previously phenotyped by two different research groups. While no ATS-related abnormalities could be discerned in either mutant by the first group at 3 months of age [23], the second group found elastic fiber proliferation in older homozygous G128E mutant mice only [22]. It is unclear if this related to the elastic fiber fragmentation seen in the vascular wall of ATS patients. Recent follow-up studies in this mouse model indicated altered redox homeostasis and mitochondrial dysfunction as pathogenic mechanisms underlying ATS [24], similar to observations in the morpholino zebrafish model [21], and in ATS knockdown cell lines (rat A10, mouse 3T3-L1) [14]. Altered redox homeostasis contributes to the pathogenesis in other aneurysm-related diseases [25].

Growing evidence that GLUT10 functions as an AA transporter has led to the hypothesis that the innate AA synthesizing capacity of mice could rescue the phenotype. Therefore, we characterized a novel mouse model for ATS, with gene disruptions of both *Slc2a10*, the causal gene for ATS, and *Gulo*, which encodes L-gunololactone, the enzyme catalyzing the last step in AA biosynthesis in rodents.

## Materials & methods

### Animals

Heterozygous knock-out mice for *Slc2a10*, in which a selection cassette replaces a sequence ranging from exon 2 to the beginning of exon 5, were purchased from Taconic Biosciences (cat. nr. TF2438), (Supplementary Figure 1a). The mutation resides in a mixed 129/SvEv-C57BL/6 genetic background. Heterozygous knock-out mice for *Gulo*, in which a selection cassette replaces exons 3 and 4, were purchased from MMRRC (RRID MMRRC_000015-UCD)(Supplementary Figure 1a). This mouse line, residing in a mixed 129/SvEv-C57BL/6 genetic background as well, was previously described by Maeda and colleagues [26]. The two lines were crossed according to a breeding scheme depicted in Supplementary Figure 1b. To allow for valid data generation and to take an influence of genetic background into account, all comparisons were made between littermates from the F2 generation. Male and female mice of the following genotypes were studied: double knockout (DKO – *Gulo*^tm1mae/tm1mae^;*Slc2a10*^−/−^), single knockout (*Gulo* KO – *Gulo*^tm1mae/tm1mae^;*Slc2a10*^+/+^ and *Slc2a10* KO – *Gulo*^+/+^;*Slc2a10*^−/−^) and wild type (WT – *Gulo*^+/+^;*Slc2a10*^+/+^). All procedures were conducted in compliance with the European Parliament Directive 2010/63/EU, and with the approval of the Ghent University ethical committee on animal experiments (Permit Number: ECD 13/17). All animals were fed ad libitum with a standard mouse breeding feed, supplemented with 300 mg/kg ascorbic acid (Ssniff^®^ Spezialdiäten) and maintained in a fully controlled animal facility (12:12 h light:dark cycle at ± 22 °C).

### Genotyping

Seven-day old mice were toe-clipped, after which genomic DNA was prepared from the tissue using the KAPA Express Extract DNA Extraction Kit (Kapa Biosystems, cat. nr. KK7103). For both *Gulo* and *Slc2a10* the DNA was PCR-amplified using specific primers detecting the wild type and mutant alleles. The genotype was determined based on PCR product presence and band size (Supplementary Table 1).

### Vascular smooth muscle cells isolation and cell culture

Mice (at an age of 25 days) were euthanized (CO_2_) and autopsied according to standard protocols. Primary VSMCs from the thoracic aorta were isolated according to the collagenase digestion method of Proudfoot and Shanahan [27]. Each cell line was derived from a single aorta. Primary VSMCs were cultured on gelatinized dishes in SmBM medium supplemented with the SmGM-2 kit (CC-3182, Lonza). Unless otherwise specified, two cell lines per genotype were assessed.

### Quantitative real-time PCR

Skin and aortic tissue from 3 male and 3 female mice of each selected genotype were harvested at the age of 9 months and submerged in RNAlater™ Stabilization Solution (Thermo Fisher Scientific, cat. nr. AM7020) prior to the RNA extraction procedure, carried out with the RNeasy^®^ Mini kit (Qiagen, cat. nr. 74106). Next, cDNA synthesis was performed using the iScript cDNA synthesis kit (Bio-Rad, cat. nr. 1708891). RT-qPCR reactions were carried out in quadruple on a LC480 machine (Roche). The reaction mix consisted of cDNA, LightCycler^®^ 1536 DNA Probes Master (Roche, cat. nr. 05502381001), LightCycler^®^ 480 High Resolution Melting Dye (Roche, cat. nr. 04909640001) and specific primers amplifying the *Slc2a10* transcript or Expressed Repeat Elements (EREs) (Supplementary Table 2) [28], to which all reactions were normalized. Data analysis was carried out with qBase+ (Biogazelle). Values plotted correspond to the means of 3 male and 3 female biological replicates. Error bars shown represent 95% confidence intervals. Data were analyzed using Welch’s ANOVA, followed by a Games-Howell post-hoc test.

### *In vivo* imaging

Serial ultrasound imaging was performed using a dedicated ultrasound apparatus (Vevo 2100, VisualSonics), equipped with a high-frequency linear array transducer (MS 550D, frequency 22–55 MHz). Imaging was carried out on mice of 6 weeks, 3, 6 and 9 months old. Ten male and 10 female mice of the selected genotypes were studied. The mice were anesthetized with 1.5% isoflurane mixed with 0.5 L/min 100% O_2_. Body temperature was maintained at 37°C, by placing the mice on a heating pad. Artery and cardiac measurements were carried out in concordance with previously established guidelines [29]. In short, the diameter measurements of the aorta at the level of the aortic root, proximal ascending aorta, distal ascending aorta, aortic arch and descending aorta were performed out, next to diameter measurements of the carotid arteries and cardiac assessment. Three cardiac cycles for each animal were analyzed (VevoLAB 1.7.0, VisualSonics). For each measurement location, a linear mixed model with covariance pattern modelling was fitted to account for correlated responses (diameters) within the same subject. Genotype, age and sex were considered as categorical explanatory variables. Starting from a saturated mean model with unstructured covariance matrix and using REML, the covariance model was simplified by comparison with simpler structures through Akaike’s Information Criterion. The following covariance matrices that were as parsimonious as possible, were selected: unstructured (aortic root), autoregressive (proximal ascending aorta), Toeplitz (distal ascending aorta), compound symmetry (all other measurement locations). Afterwards the fixed part of the model was simplified by testing simpler models using ML. All final models included the main effects of genotype, age, and sex without interaction terms and were refitted with REML. Values plotted correspond to the means of ±10 biological replicates. Error bars shown represent 95% confidence intervals.

### Vascular corrosion casting

The vascular corrosion casting technique, based on the injection of a polymer to capture the 3D structure of the vasculature, was performed on 3 male and 3 female mice of each selected genotype at the age of 9 months as previously described [23]. Briefly, 2 to 3 ml Batson solution (Omnilabo International, Batson’s #17 corrosion kit) was injected in the abdominal aorta through a 26G catheter. After completion of the polymerization reaction, mouse bodies were macerated overnight in a 25% KOH solution. The resulting casts were cleaned, evaluated and photographed using a dissecting microscope, equipped with a 5-megapixel camera (Leica).

### Histology

Skin (9 months), aorta (9 months) and eye (3 weeks) tissue was collected from 3 male and 3 female mice of each selected genotype for histological analysis. Samples were formalin-fixed and paraffin-embedded, after which 5 μm-thick paraffin sections were made. Sections were subjected to the standard hematoxylin & eosin, Verhoeff-Van Gieson and picrosirius red histological staining procedures, prior to visualization on a Zeiss Axio Observer Z1 microscope.

### Transmission Electron Microscopy

Sample fixation was carried out in a 4% formaldehyde (EM grade), 2.5% glutaraldehyde (EM grade), 0.1M cacodylate buffer solution. Fixed samples were stained en bloc with OsO4 and uranyl acetate, followed by dehydration and embedding in Epon, as previously described [30]. Thin sections (60 nm) were placed on formvar-coated grids, and subsequently counterstained with 7% methanolic uranyl acetate and lead citrate. Imaging was carried out with a Tecnai 12 transmission electron microscope at 120 kV. Collagen diameter measurements were carried out with Fiji [31]. Error bars shown represent 95% confidence intervals. Data were analyzed with a two-tailed t-test.

### Immunofluorescence of extracellular matrix

ECM protein production by VSMCs was determined by immunofluorescence. VSMCs were seeded at 50,000 cells/well in 8-well removable chamber slides and grown for 7 days to allow ECM deposition. VSMCs were fixed with an ice-cold 70:30 methanol:acetone mixture for 5 minutes and washed with PBS. Coverslips were blocked for 1 hour with PBS supplemented with 10% normal goat serum (X0907, Agilent). Primary antibodies (Supplementary Table 3) were incubated overnight at 4°C in PBS. Coverslips were washed three times with PBS for 5 minutes each prior to incubation with the secondary antibody for 1.5 hours at room temperature (1:1,000, anti-rabbit Alexa Fluor 594, Molecular Probes). Coverslips were washed and mounted to glass slides with Vectashield supplemented with DAPI (H-1200, Vector laboratories) and sealed with nail polish. Images were recorded on a wide field epifluorescent microscope (Axio Imager D2, Zeiss). Quantification of the immunofluorescent signal was performed by calculating the corrected total cell fluorescence (CTCF) of the ECM components corrected for the number of nuclei. The CTCF was determined by setting a color threshold to select the fibers in the image with Fiji image analyzing software [31] and determining the integrated density of this area (intensity of the fluorescence). This measurement was corrected for the background fluorescence and the total area of the fibers and results in the CTCF. The CTCF was then divided by the number of nuclei that were present in the measured image. Data were corrected for outliers with the Grubbs’ test for outliers [32]. Statistical analysis was performed with a non-parametric Mann-Whitney test. Significance was tested 2-tailed. Results are expressed as mean ± SD.

### TGFβ stimulation

VSMCs were seeded in 6-well plates to reach confluence and were allowed to attach for 24 hours. The following day, medium was changed to SmGM-2 medium without FCS and VSMCs were serum deprived for 24 hours prior to TGFβ stimulation. Protein samples were collected after 0 minutes, 15 minutes, 30 minutes, 1 hour and 4 hours of stimulation with TGFβ1 (4342-5, Biovision). Cells were scraped in PBS supplemented with protease inhibitor cocktail (1:100, 11836145001, Roche applied science) and phosphatase inhibitor cocktail (1:100, P0044, Sigma) and lysed in an equal volume of 2x Laemmli buffer (4% SDS, 20% glycerol, 120mM Tris pH 6,8) supplemented with protease inhibitor cocktail and phosphatase inhibitor. Lysates were first cleared from large DNA by passing through a 25G needle and then heated to 65°C for 10 minutes. Protein concentrations were measured with the Lowry protein assay as described [33]. Equal amounts of protein were separated for size by SDS-PAGE and then transferred to a PVDF membrane (1 hour, 100V, Immobilon) and blocked with either 3% milk in PBS supplemented with 0.1% Tween-20 (1 hour, room temperature). The primary antibody was incubated for 45 minutes at room temperature or overnight at 4°C for phosphorylated Smad2 (see Supplementary Table 3 for primary antibodies). The membranes were washed 5 times with 0.1% Tween-20 in PBS and then incubated with horseradish peroxidase-conjugated secondary antibodies (1:2,000, Jackson ImmunoResearch) for 1 hour at room temperature. Bound secondary antibodies were detected with an Amersham Imager 600 (GE Healthcare Life Sciences) using chemiluminescence.

Band intensity was quantified using Fiji image analyzing software [31]. Data were corrected for outliers with the Grubbs’ test for outliers [32]. Statistical analysis was performed with a non-parametric Mann-Whitney test. Significance was tested 2-tailed. Results are expressed as mean ± SD.

### Mitochondrial respiration

Oxygen consumption rate (OCR) was measured using an XF-24 Extracellular Flux Analyzer (Seahorse Bioscience). Respiration was measured in XF assay media (non-buffered DMEM), in basal conditions and in response to 1 μM oligomycin (ATP synthase inhibitor), 1 μM fluoro-carbonyl cyanide phenylhydrazone (FCCP, uncoupler), 1 μM antimycin A (complex III inhibitor). Smooth muscle cells were seeded at a density of 30,000 cells/well and analyzed after 24 hours. Optimal cell densities were determined experimentally to ensure a proportional response to FCCP with cell number. For these experiments 6-8 wells were measured per time point [34, 35]. Data were corrected for outliers with the Grubbs’ test for outliers [32]. Statistical analysis was performed with a non-parametric Mann-Whitney test. Significance was tested 2-tailed. Results are expressed as mean ± SD.

## Results

### Generation of the *Slc2a10*;*Gulo* mouse model

We developed a novel mouse model for ATS in a mixed 129/SvEv-C57BL/6 genetic background by crossing constitutive knock-out lines for *Gulo* and *Slc2a10*, both harboring a selection cassette replacing multiple exons (Supplementary Figure 1a). In order to take a possible effect of a variable genetic background on the acquired data into account, all studies were performed on littermates from the F2 generation (Supplementary Figure 1b). RT-qPCR-based expression analysis, using primers targeting the sequence replaced by the selection cassette, confirms the absence of wild type *Slc2a10* RNA in skin and aorta obtained from *Slc2a10*^−/−^ animals (DKO and *Slc2a10* KO) (Figure 1). Double knockout mice (DKO – *Gulo*^tm1mae/ tm1mae^;*Slc2a10*^−/−^) are born with expected Mendelian frequency and have a normal lifespan to at least 9 months of age, similar to their control littermates (*Gulo* KO – *Gulo*^tm1mae/ tm1mae^;*Slc2a10*^+/+^, *Slc2a10* KO – *Gulo*^+/+^;*Slc2a10*^−/−^ and WT – *Gulo*^+/+^;*Slc2a10*^+/+^). Double knock-out mice develop normally, and show no obvious gross anomalies, skeletal dysmorphisms or skin abnormalities. However, the weight of female DKO mice is slightly reduced, at each measurement time point (Supplementary Figure 2).

**Figure 1:**
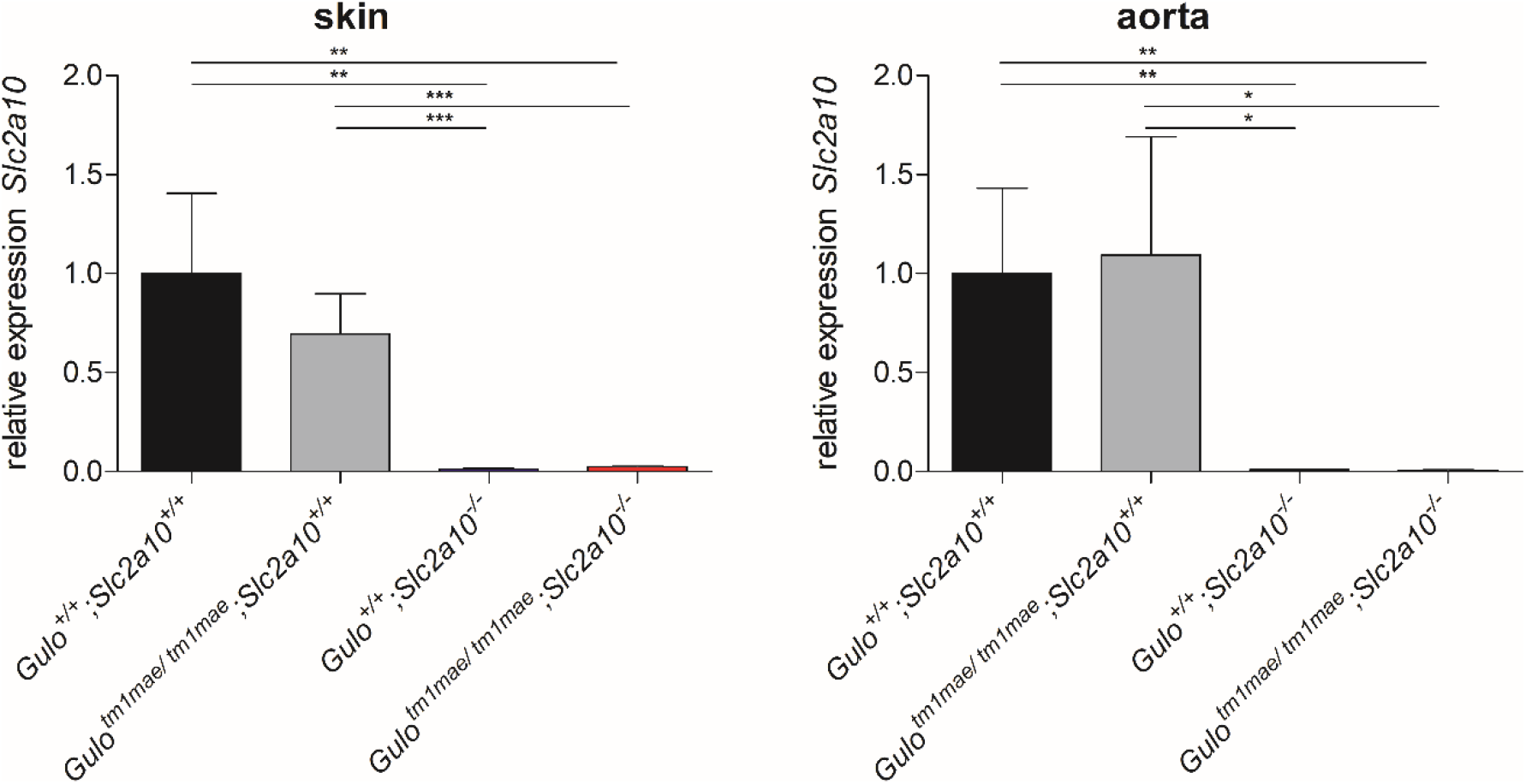
*Slc2a10* expression levels in skin and aortic tissue. RT-qPCR analysis identifies a near absence of *Slc2a10* WT mRNA in mice homozygous for *Slc2a10* mutations (*Gulo*^+/+^;*Slc2a10*^−/−^ and *Gulo*^tm1mae/ tm1mae^;*Slc2a10*^−/−^), compared to mice harboring the *Slc2a10* wild type allele (*Gulo*^tm1mae/tm1mae^;*Slc2a10*^+/+^ and *Gulo*^+/+^;*Slc2a10*^+/+^). Displayed average values represent the relative expression data in skin or aorta for 3 male and 3 female mice. Error bars represent the SD, *: p≤0,05; **: p≤0,01; ***: p≤0,001.

### *Slc2a10;Gulo* double knock-out mice do not copy the human ATS phenotype completely

We carried out serial ultrasound measurements at multiple locations in the cardiovascular system of 10 male and 10 female mice of each selected genotype (DKO, *Slc2a10* KO, *Gulo* KO, WT) at 6 weeks, 3, 6, and 9 months of age. Diameters of the ascending aorta, distal ascending aorta, aortic arch and descending aorta show a subtle but significant decrease in DKO mice compared to their wild type littermates, for a given age and sex (Figure 2). More distal measurements (carotid arteries) do not reveal any significant diameter alterations (Supplementary Figure 3). Fractional shortening and ejection fraction are comparable between the selected genotypes at 6 weeks and 3 months of age (Supplementary Figure 4). We did not detect aortic or mitral valve insufficiency using pulsed Doppler (data not shown). Detailed microscopic analysis of the aortic tree, eye arterioles and the circle of Willis, obtained from polymer replicas of the cardiovascular system at 9 months of age, does not indicate any omnipresent structural anomalies such as tortuosity, abnormal implantation of the aortic side-branches or aneurysms. However, one out of three analyzed female DKO mice has a local stenosis in the vertebral artery (Figure 3, Supplementary Figure 5). At 9 months, elastic fiber staining of aortic tissue shows mild age-related changes, such as sporadic elastic fiber fragmentation or accumulation, in the *tunica media* of the aortic wall across all selected genotypes.

**Figure 2:**
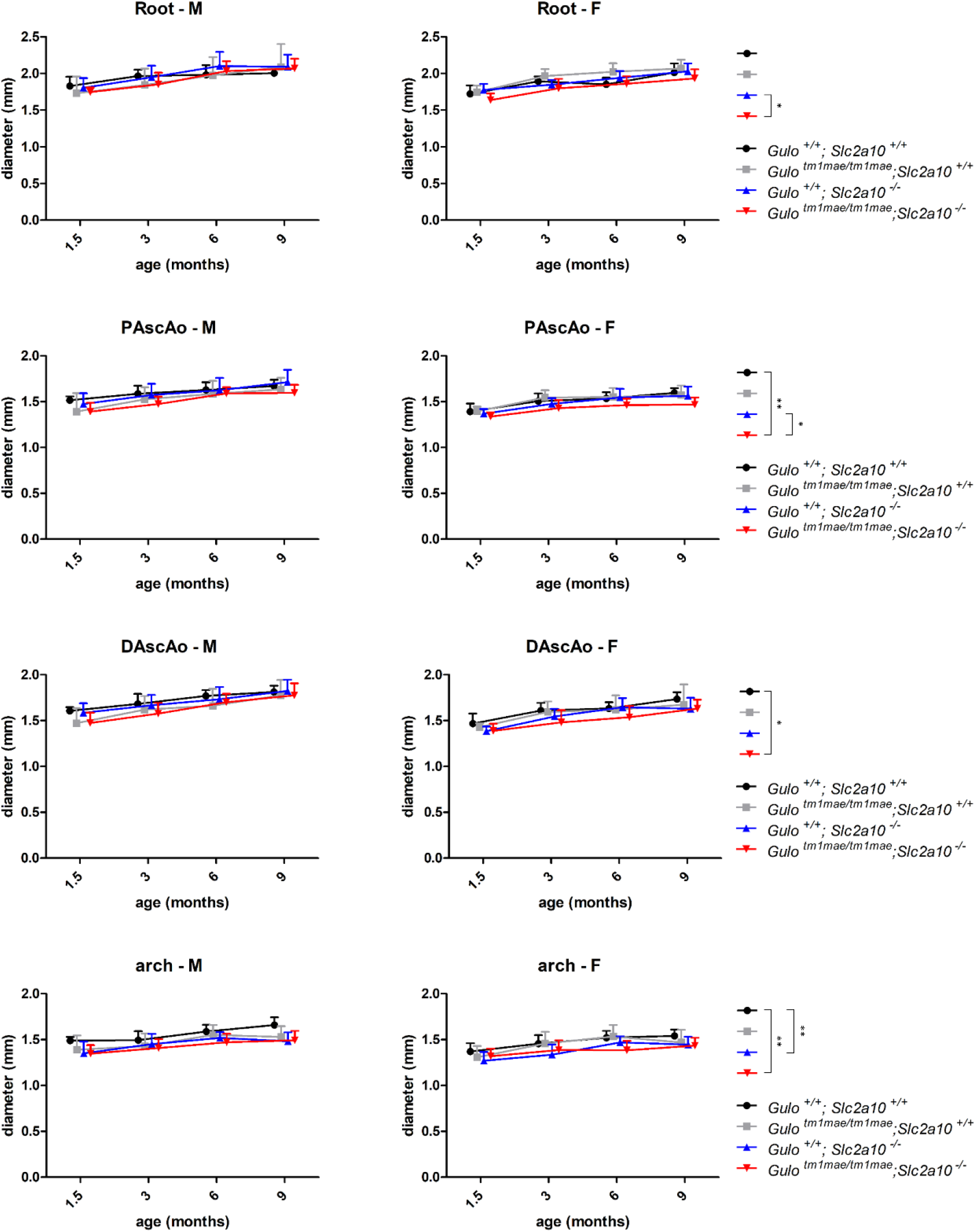
Serial ultrasound measurements obtained in *Gulo*^+/+^;*Slc2a10*^+/+^, *Gulo*^tm1mae/tm1mae^;*Slc2a10*^+/+^, *Gulo*^+/+^;*Slc2a10*^−/−^ and *Gulo*^tm1mae/tm1mae^;*Slc2a10*^−/−^ mice at 6 weeks, 3, 6 and 9 months of age. Shown data was acquired at the level of the aortic root, proximal ascending aorta, distal ascending aorta and aortic arch. Left and right column images represent data obtained in male (M) and female (F) animals respectively. Shown p-values were obtained using a linear mixed model with covariance pattern modeling, without interactions. As a result, p-values should be interpreted when making the comparison for a given age and sex. Error bars shown represent 95% confidence intervals. *: p ≤ 0,05; **: p ≤ 0,01. PAscAo (proximal ascending aorta), DAscAo (distal ascending aorta).

**Figure 3:**
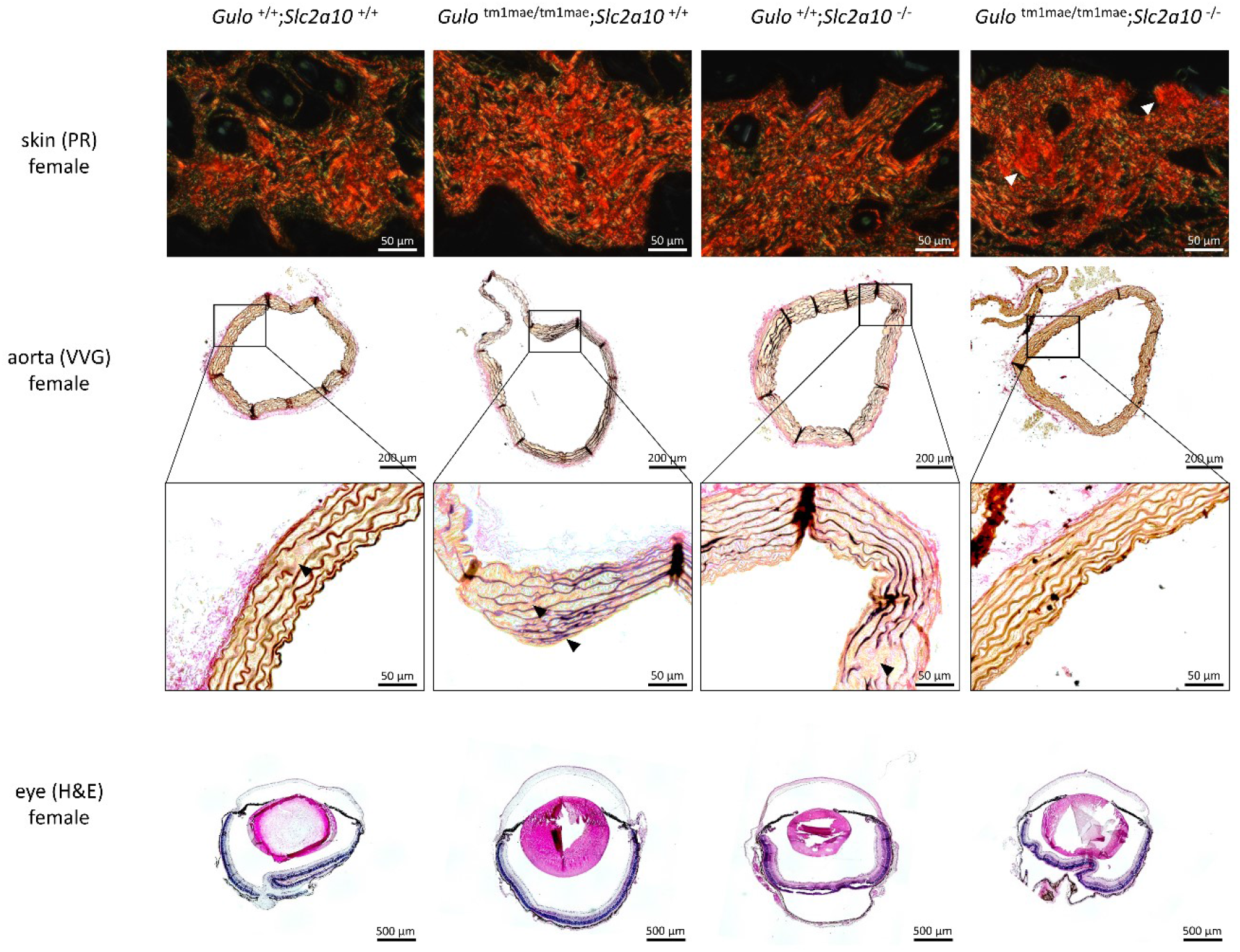
Vascular corrosion casts of *Gulo*^+/+^;*Slc2a10*^+/+^, *Gulo*^tm1mae/tm1mae^;*Slc2a10*^+/+^, *Gulo*^+/+^;*Slc2a10*^−/−^ and *Gulo*^tm1mae/ tm1mae^;*Slc2a10*^−/−^ female mice. Representative images of the aortic arch and its side branches, the circle of Willis and the eye arterioles are shown. A local narrowing in the vertebral artery in the circle of Willis could be observed in one *Gulo*^tm1mae/ tm1mae^;*Slc2a10*^−/−^ mouse (white-framed enlargement). Scale bars: 2 mm.

**Figure 4:**
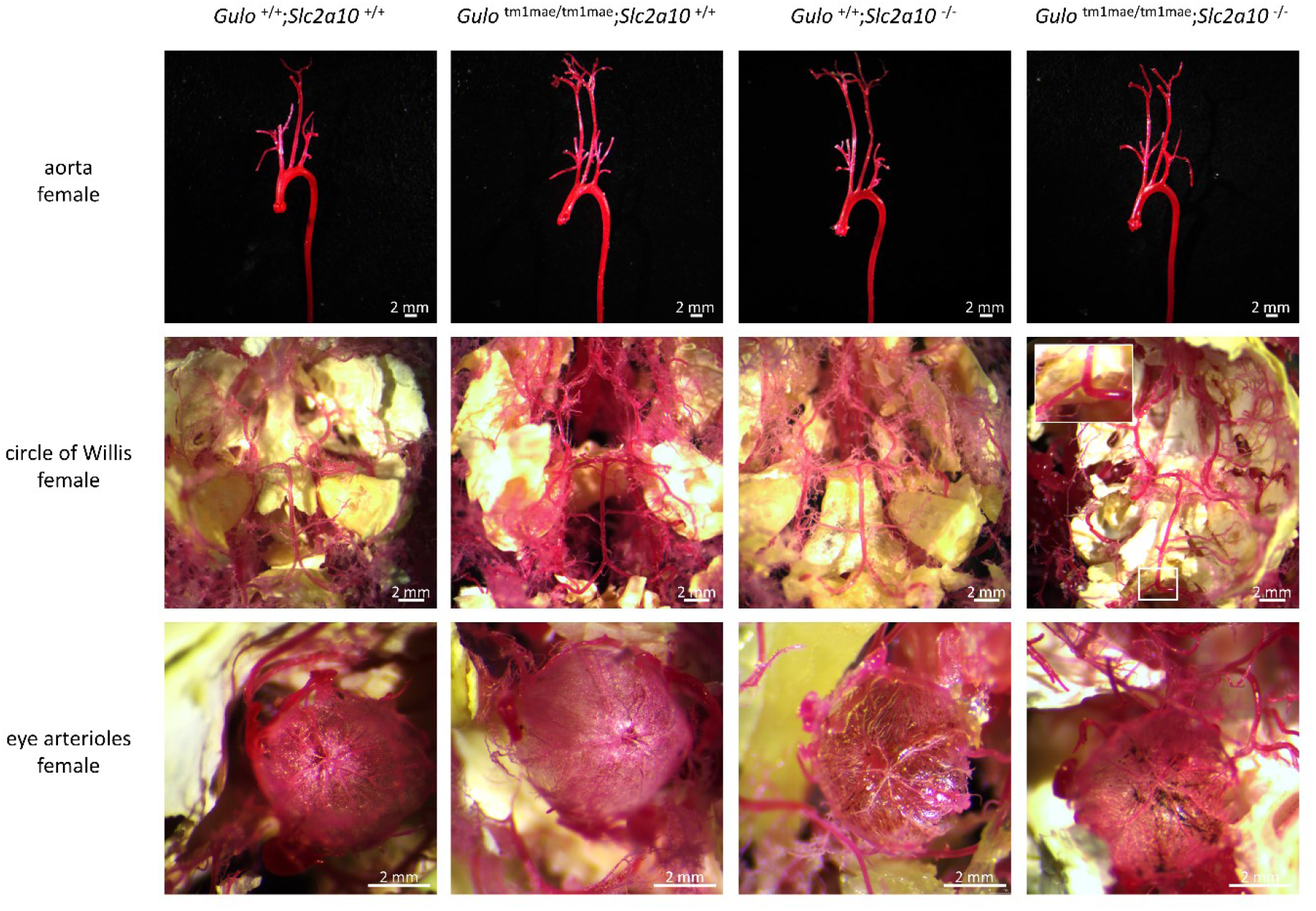
Histological analysis of *Gulo*^+/+^;*Slc2a10*^+/+^, *Gulo*^tm1mae/tm1mae^;*Slc2a10*^+/+^, *Gulo*^+/+^;*Slc2a10*^−/^and *Gulo*^tm1mae/ tm1mae^;*Slc2a10*^−/−^ female mice. Picrosirius red (PR) polarization staining (upper row) for collagen reveals the presence of patches with accumulated collagen deposition (white arrows) in the skin of *Gulo*^tm1mae/ tm1mae^;*Slc2a10*^−/−^ mice. Verhoeff-Van Gieson (VVG) elastic fiber staining (middle row) shows mild elastic fiber anomalies in the aortic wall of all studied mice (black arrows). Hematoxylin & eosin (H&E) staining (bottom row) in eye tissue does not reveal major structural abnormalities in the cornea.

Picrosirius red polarization staining on skin tissue shows no major alterations in collagen deposition, but patches of increased collagen deposition can be discerned in female DKO mice (Figure 4, Supplementary Figure 6). DKO mice do not show keratoconus or keratoglobus (Figure 4) [1].

Ultrastructural Transmission Electron Microscopy (TEM) analysis of female skin samples shows a normal elastin fiber architecture. Irregularities in collagen diameter are significantly different between DKO and WT mice (Figure 5).

**Figure 5:**
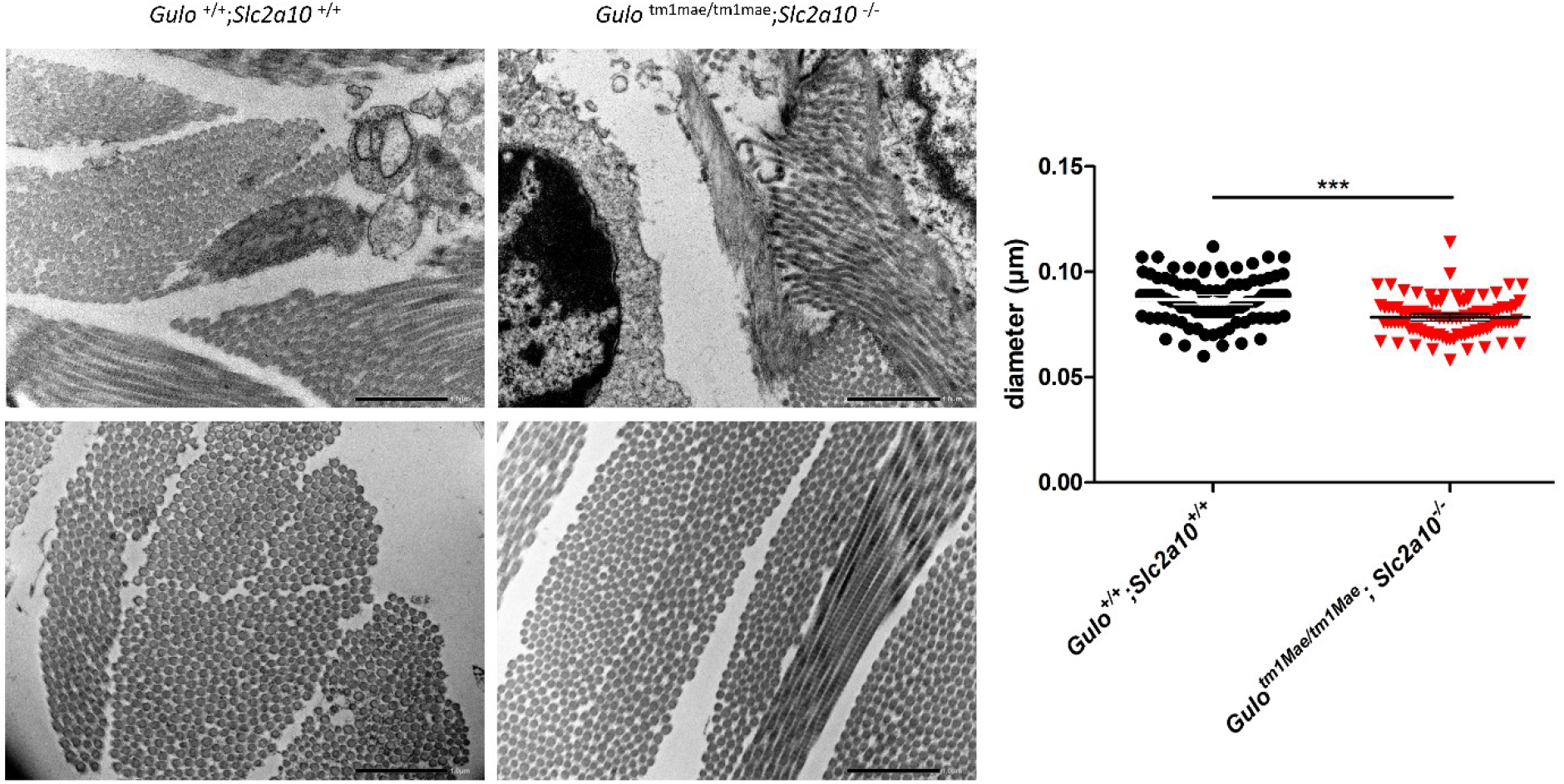
Electron microscopic findings in skin samples, and collagen diameter quantification. *Gulo*^tm1mae/tm1mae^;*Slc2a10*^−/−^ mice show decreased collagen fiber diameters, compared to *Gulo*^+/+^;*Slc2a10*^+/+^ mice. Scale bar: 10 μm, Error bars shown represent 95% confidence intervals. ***: p ≤ 0,001.

### *In vitro* analysis of *Slc2a10;Gulo* double knock-out VSMCs shows decreased ECM deposition

We examined the ECM deposition in VSMCs of the DKO and WT genotypes (Figure 6). In contrast to WT VSMCs, DKO VSMCs do not show a base network of recognizable and organized fibronectin fibers stretching between the cells, but present with a reduced intensity of fibronectin staining with a more fuzzy appearance (Figure 6A, Supplementary Figure 7). A similar observation is made for Fibrillin-1 where stained fibers appeared thinner in DKO VSMCs (Figure 6B, Supplementary Figure 7). WT VSMCs show clear staining of tropoelastin that appears in a fibrillar structure (Figure 6C), next to the presence of a strong fiber network of both fibulin-4 (Figure 6D) and fibulin-5 (Figure 6E). In DKO VSMCs, a reduced staining of fibulin-4 and fibulin-5 (Figure 6D and E, Supplementary Figure 7) and an overlapping network of tropoelastin and fibulins (Figure 6C, Supplementary Figure 7) can be observed. We further assessed LTBP-4, which is present in clear fibrillar structures (Figure 6F) in both WT and DKO VSMCs, but appears less intense in DKO VSMCs, organized in a more densely packed fibril network (Figure 6F, Supplementary Figure 7). Taken together, this data shows that DKO VSMCs display decreased deposition of the various components of the ECM compared to WT VSMCs.

**Figure 6:**
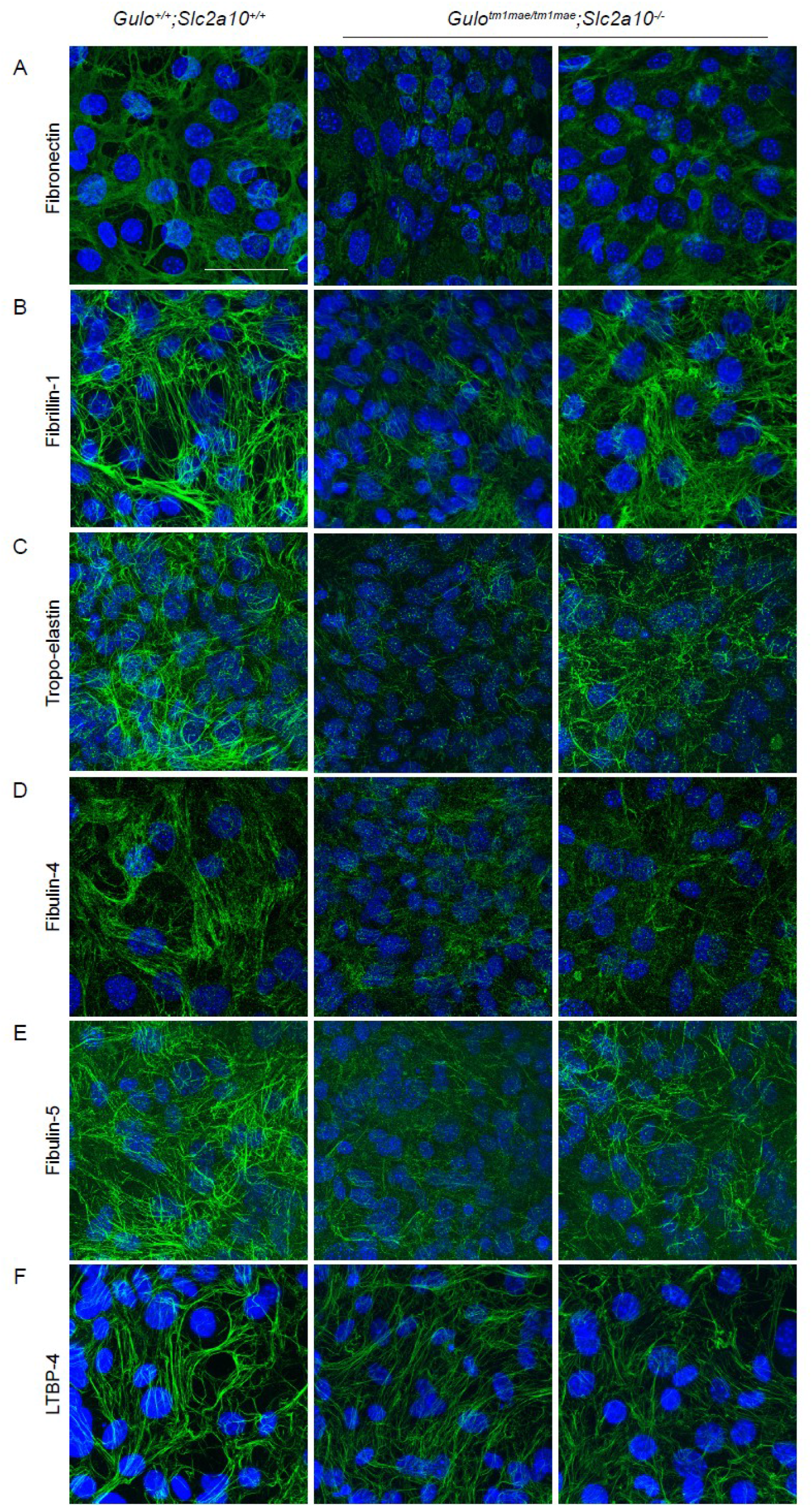
*Slc2a10;Gulo* double knock-out VSMCs show decreased ECM deposition. Immunofluorescent images show deposition of multiple ECM components after 7 days of culture. *Gulo*^tm1mae/tm1mae^;*Slc2a10*^−/−^ VSMCs show less distinct ECM structures compared to *Gulo*^+/+^;*Slc2a10*^+/+^ VSMCs. Scale bars represent 100um.

### *Slc2a10;Gulo* double knock-out VSMCs do not show altered activation of canonical TGFβ signaling

We investigated canonical TGFβ signaling activation in VSMCs derived from WT and DKO mouse aortas. Under an unstimulated serum deprived condition, DKO VSMCs show similar levels of Smad2 phosphorylation compared to WT VSMCs (Figure 7A and B). Following TGFβ stimulation, both WT and DKO VSMCs demonstrate a strong and comparable increase in phosphorylation of Smad2 after 15 minutes (Figure 7A). Quantification of phosphorylated Smad2 to total Smad2 ratio reveals a further increase to its maximum at 60 minutes of stimulation after which it decreases (Figure 7A and B). In conclusion, our data do not support aberrations in the activation of the canonical TGFβ signaling pathway in DKO VSMCs compared to control VSMCs.

**Figure 7:**
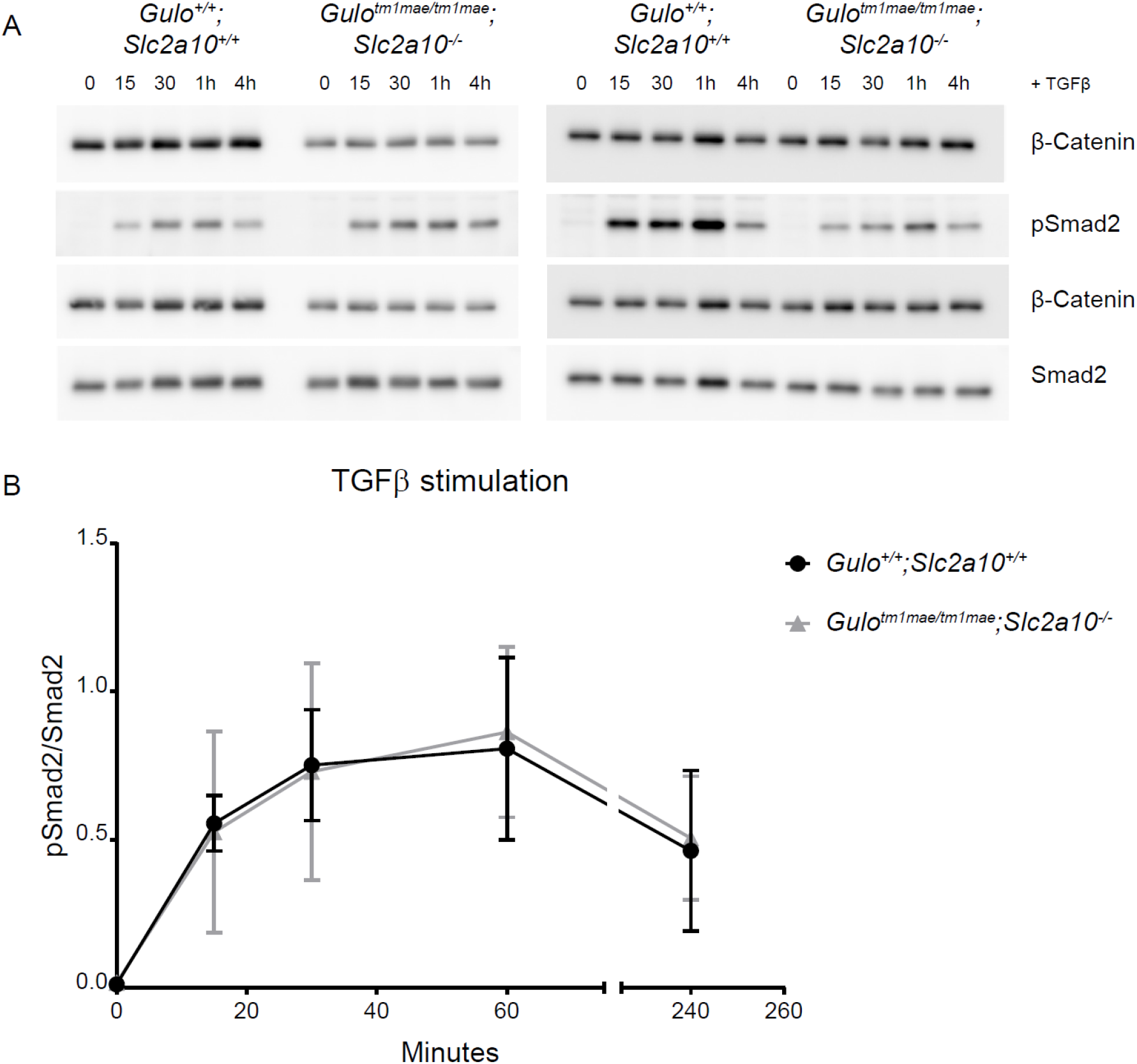
*Slc2a10;Gulo* double knock-out VSMCs do not show aberration in activation of the canonical TGFβ signaling pathway. *Gulo*^tm1mae/tm1mae^;*Slc2a10*^−/−^ VSMCs do not show increased phosphorylation of Smad2 without TGFβ stimulation and do not differ from *Gulo*^+/+^;*Slc2a10*^+/+^ VSMCs in basal conditions. Stimulation with TGFβ led to increased phosphorylation of Smad2 in both *Gulo*^+/+^;*Slc2a10*^+/+^ and *Gulo*^tm1mae/tm1mae^;*Slc2a10*^−/−^ VSMCs. There was no difference in pSmad2/Smad2 ratio between the two genotypes. Data shows 2 independent cell lines from 2 experiments, data point represent average values of an independent cell line per experiment. Results are depicted as mean ±SD.

### *Slc2a10;Gulo* double knock-out VSMCs show a reduced maximum OCR

We analyzed the oxygen consumption rate (OCR) of DKO compared to WT VSMCs (Figure 8A). Phase I represents the basal and unstimulated OCR. Addition of oligomycin in phase II inhibits the ATP synthase and reduces the OCR. In Phase III, treatment with fluoro-carbonyl cyanide phenylhydrazone (FCCP) uncouples oxygen consumption from ATP production and raises OCR to its maximum. Addition of antimycin A in phase IV inhibits complex III of the mitochondria and blocks mitochondrial respiration. Figure 8B and C contain the quantification of the mean value of phase I and III per genotype. Basal OCR does not differ between both genotypes (Figure 8A and B). However, when VSMCs are treated with FCCP to reach maximum respiration, DKO VSMCs display decreased maximum respiration rates compared to WT VSMCs (Figure 8A and C).

**Figure 8:**
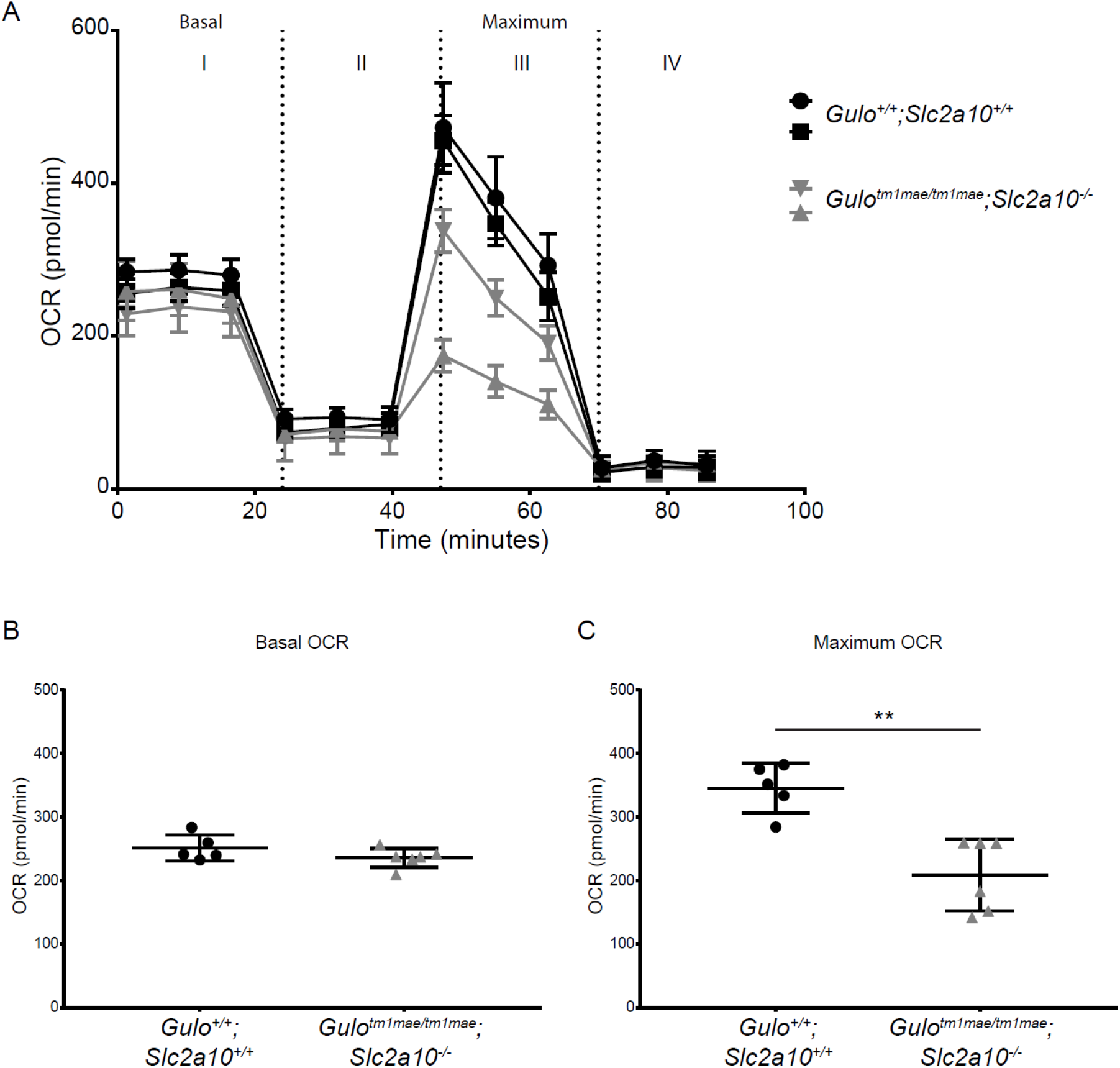
*Slc2a10;Gulo* double knock-out VSMCs show a reduced maximum OCR. In basal conditions, *Gulo*^tm1mae/tm1mae^;*Slc2a10*^−/−^ and *Gulo*^+/+^;*Slc2a10*^+/+^ VSMCs do not differ in OCR, while treatment with FCCP results in a decreased maximum OCR of *Gulo*^tm1mae/tm1mae^;*Slc2a10*^−/−^ VSMCs. Data shows 2 independent cell lines from 3 experiments, data point represent average values of an independent cell line per experiment. Results are depicted as mean ±SD. ** p<0,01.

## Discussion

Arterial tortuosity may occur in both innate and acquired vasculopathies, such as persistent hypertension and neoangiogenesis, and may associate with an increased vascular risk, including aneurysm formation and rupture. Monogenic conditions may be helpful in delineating the underlying pathophysiology of altered vascular patterning. In this study, we developed and characterized a novel mouse model for ATS, since the previously reported mouse models failed to recapitulate the major hallmarks of the human disease [22, 23]. Our model addressed two concerns regarding the previous models. Firstly, we developed a *Slc2a10* knock-out, which abolishes GLUT10 function while the previously reported missense mutations were only *predicted* to be pathogenic [22, 23]. Secondly, we impaired AA synthesis in our model by disrupting the *Gulo* gene, aiming to mimic the physiological conditions in human more reliably. Indeed, *Gulo* encodes L-gulonolactone oxidase, an endoplasmic reticulum membrane bound enzyme that catalyzes the last step in the vitamin C biosynthesis pathway [36]. It became mutated in many species, such as teleost fishes, guinea pigs and anthropoid primates, including humans, making these species unable to synthesize vitamin C [37]. Vitamin C plays an essential role as an anti-oxidant and in collagen synthesis. Since recent evidence indicates that GLUT10 function may include transport of DHA into subcellular compartments [14, 17, 24], it could be hypothesized that murine developmental processes do not suffer from *Slc2a10* mutations and localized vitamin C hypovitaminosis, which would explain the absence of a relevant phenotype in *Slc2a10* mutant mice [16, 23].

Nevertheless, we only observed mild cardiovascular manifestations in the double knock-out ATS mouse model. Ultrasound-guided characterization revealed a subtle stenosis from the proximal ascending aorta to the descending aorta in DKO mice, but only statistically significant when compared with their wild type littermates. Moreover, the reduced weight in the double knock-out mice further questions the biological relevance of these results.

Histopathology of ATS patient vascular tissue invariably shows substantial fragmentation of the elastic laminae in the *tunica media* [1] while conversely the previously described Glut10^G128E^ mouse model mainly presented with increased arterial wall thickness, most prominent at the age of 15 months [22, 24]. Our model only reveals mild elastic fiber anomalies in all genotypes, that are likely age-related changes. Nevertheless, dermal collagen histology revealed patches of increased collagen deposition in female DKO mice, which has been observed in skin tissue of ATS patients [1], and might contribute to their ‘doughy’ skin characteristics. Ultrastructural analysis of skin tissue of double knockout mice additionally revealed collagen fiber diameter alterations in DKO mice, but did not reveal major defective elastin deposition, which is in contrast to TEM observations in ATS patients [1].

The mild phenotypic abnormalities observed seem to be more pronounced in female mice, suggesting a sex discrepancy, even for nulliparous mice. Influences of sex in connective tissue homeostasis and disease have been previously recognized. However, most studies indicate that male mice models for connective tissue disorders show more severe phenotypes than female mice [38–41], which is in contrast to our study.

Despite the absence of a major clinical or histological phenotypes, we did identify defective ECM deposition in cultured VSMCs of DKO mice. We observed abnormalities in the fibronectin network that serves as a base network for ECM assembly, and in fibrillin-1 microfibrillar scaffolds that form the structural basis for elastic fibers. Moreover, VSMC from DKO mice show reduced fibulin-4, −5 and LTBP4 staining, and fail to establish an elastin network. These results are in line with findings by Zoppi *et al.*, who previously showed ECM disarray in skin fibroblasts of ATS patients [12]. Also, a recent study revealed an abnormal microfibrillar scaffold and incomplete elastic core assembly in skin samples of ATS patients [1], that likely reflect defective ECM assembly at multiple levels, as shown in our model. Since defective ECM assembly is often associated with vascular phenotypes [42, 43], this might indicate that mice may dispose of other mechanisms that overcome the deleterious defects of abnormal ECM assembly.

In several aneurysm syndromes, dysregulation of the canonical TGFβ signaling pathway is at play in aneurysm formation [45, 46]. However, while we observed ECM abnormalities, DKO VSMCs do not show altered phosphorylation of Smad2 upon TGFβ stimulation over time as well as under basal, unstimulated, conditions. An initial report indicated a prolonged signal increase for downstream readouts of TGFβ signalling in arterial tissue of (one) ATS patient [3]. This finding remained unconfirmed on both fibroblast cultures [12], skin or artery tissue [1] in an extended number of patients. A role for non-canonical αvβ3 integrin-mediated TGFβ signalling in ATS pathomechanisms has been suggested [12]. Our results add to the uncertainty regarding the role of TGFβ signalling in the pathomechanisms underlying many elastic fiber diseases, but cannot account for a possible contribution of spatiotemporal regulation of TGFβ signalling.

A number of studies have pointed towards a key role for oxidative stress in ATS pathogenesis. For instance, GLUT10 has been identified as a mitochondrial DHA transporter, and it was shown that increased mitochondrial targeting of GLUT10 and associated increased mitochondrial DHA uptake was triggered by stress and aging conditions [14, 24]. In a morpholino *slc2a10* knockdown zebrafish model, altered OCR and differential gene expression analysis showed involvement in oxidative signalling [21]. In the Glut10^G128E^ mouse model, Syu and coworkers similarly identified a decreased maximum OCR in early passage Glut10^G128E^ VSMCs [24]. In this study, we compared basal and maximal OCR in VSMCs of double knock-out and control mice and identified a normal basal OCR in DKO VSMCs. However, following treatment with FCCP, a decreased maximum OCR could be observed in DKO VSMCs, compared to WT. Since basal OCR was comparable in DKO and WT VSMCs, the need for a stressor to dysregulate mitochondrial function in DKO VSMCs could explain the reduced maximum OCR after FCCP treatment. However, it has been suggested that an aberrant maximal OCR only corresponds to the initial symptoms of mitochondrial dysfunction caused by *SLC2A10* mutations, due to ROS that trigger arterial wall remodeling. In its turn, this leads to additional ROS production, and disease progression promotes further decline in mitochondrial function. It is also plausible that mitochondrial dysfunction is a more ubiquitous phenomenon associated with aneurysmal disease. Van der Pluijm *et al.* investigated mitochondrial function in Fibulin-4^R/R^ and Tgfbr-1^M318R/+^ mouse VSMCs and found decreased mitochondrial respiration in these mouse models. In addition, skin fibroblasts of aneurysmal patients with *FBN1, TGFBR2* or *SMAD3* mutations revealed mitochondrial dysfunction as well [25].

Finally, due to the mild nature and low abundance of relevant phenotypic alterations, it could be argued that this novel ATS double knock-out mouse model does not appear to be a suitable disease model to study the pathomechanisms underlying ATS [22, 23]. While the lack of a severe phenotype could be attributed to the genetic background of the used mouse strains, possibly hindering phenotypic penetrance [47], the underlying cause could also be biological robustness. This is a phenomenon by which the functionality of the mutated gene is maintained by increased expression of related genes [48, 49], as was demonstrated earlier in a number of other mouse disease models [50, 51]. One mechanism of biological robustness is transcriptional adaptation [52, 53]. This mechanism is triggered by mRNA degradation products and is believed to act via sequence similarity [54, 55]. Since *Slc2a10* is a member of a large family of facilitative glucose transporter proteins harboring a high sequence similarity, this mechanism is plausible [56].

Nevertheless, the current double knock-out mouse model could be of importance in revealing redundant gene networks, alternative pathways or modifier genes, possibly providing clues for future therapies. Moreover, since isolated VSMCs of double knock-out mice mimicked cellular disturbances found in cell cultures of ATS patients, these VSMCs could potentially be used for further identification of the molecular mechanisms that underlie ATS, while analyses at the histological level may reveal adaptive mechanisms that contribute to the phenotypic rescue.

In conclusion, we developed a novel mouse model for ATS, double-knockout for *Gulo* and *Slc2a10*. While our model does not phenocopy human ATS, it did reveal alterations at the cellular level including disturbed ECM assembly and altered cellular respiration. Our model confirms a role for ascorbate synthesis and compartmentalization in the disease mechanisms leading to ATS and may be helpful to identify the mechanisms underlying the phenotypic rescue in mice.

## Author contributions

Conceptualization: AnB, MR, BC, AW, PC, IvdP; Methodology: AnB, MV, MR, AW, JB, IvdP, JE, BC; Data acquisition: AnB, AuB, MV, MR, CC, JB, SB; Data analysis: AnB, JB; Contributed reagents/materials/analysis tools: DR, BD, CV; Writing – original draft: AnB, JB, AW. Writing – review: all authors.

## Acknowledgements

This work was supported by Ghent University (Methusalem BOFMET2015000401 to Anne De Paepe) and by Research Foundation – Flanders (FWO, FWOOPR2013025301). B.C. is a Senior Clinical Investigator of the Research Foundation – Flanders (FWO). M.R. was supported by the Research Foundation – Flanders (FWO) as a postdoctoral fellow. Ghent University Hospital is a member of the European Reference Networks for vascular and skin disorders (VASCERN and ERN-Skin).

