## Supplementary data for "SLC2A knockout mice deficient in ascorbic acid synthesis recapitulate aspects of arterial tortuosity syndrome and display mitochondrial respiration defects"

---

Annekatrien Boel [1], Joyce Burger [2], Marine Vanhomwegen [1], Aude Beyens [1], Marjolijn Renard [1], Sander Barnhoorn [2], Christophe Casteleyn [3], Dieter P. Reinhardt[4], Benedicte Descamps [5], Christian Vanhove [5], Ingrid van der Pluijm [2,6], Paul Coucke [1], Andy Willaert [1], Jeroen Essers [2,6,7], Bert Callewaert [1]

[1] Center For Medical Genetics Ghent, Department of biomolecular medicine, Ghent University, Ghent, Belgium

[2] Department of Molecular Genetics, Erasmus University Medical Center, Rotterdam, the Netherlands; Department of Clinical Genetics, Erasmus University Medical Center, Rotterdam, the Netherlands

[3] Department of Morphology, Faculty of Veterinary Medicine, Ghent University, Merelbeke, Belgium

[4] McGill University, Faculty of Medicine, Department of Anatomy and Cell Biology and Faculty of Dentistry, Montreal, Quebec, Canada

[5] Infinity (IBiTech-MEDISIP), Department of Electronics and Information Systems, Ghent University, Ghent, Belgium

[6] Department of Vascular Surgery, Erasmus University Medical Center, Rotterdam, the Netherlands

[7] Department of Radiation Oncology, Erasmus University Medical Center, Rotterdam, the Netherlands

---

### Supplementary figures

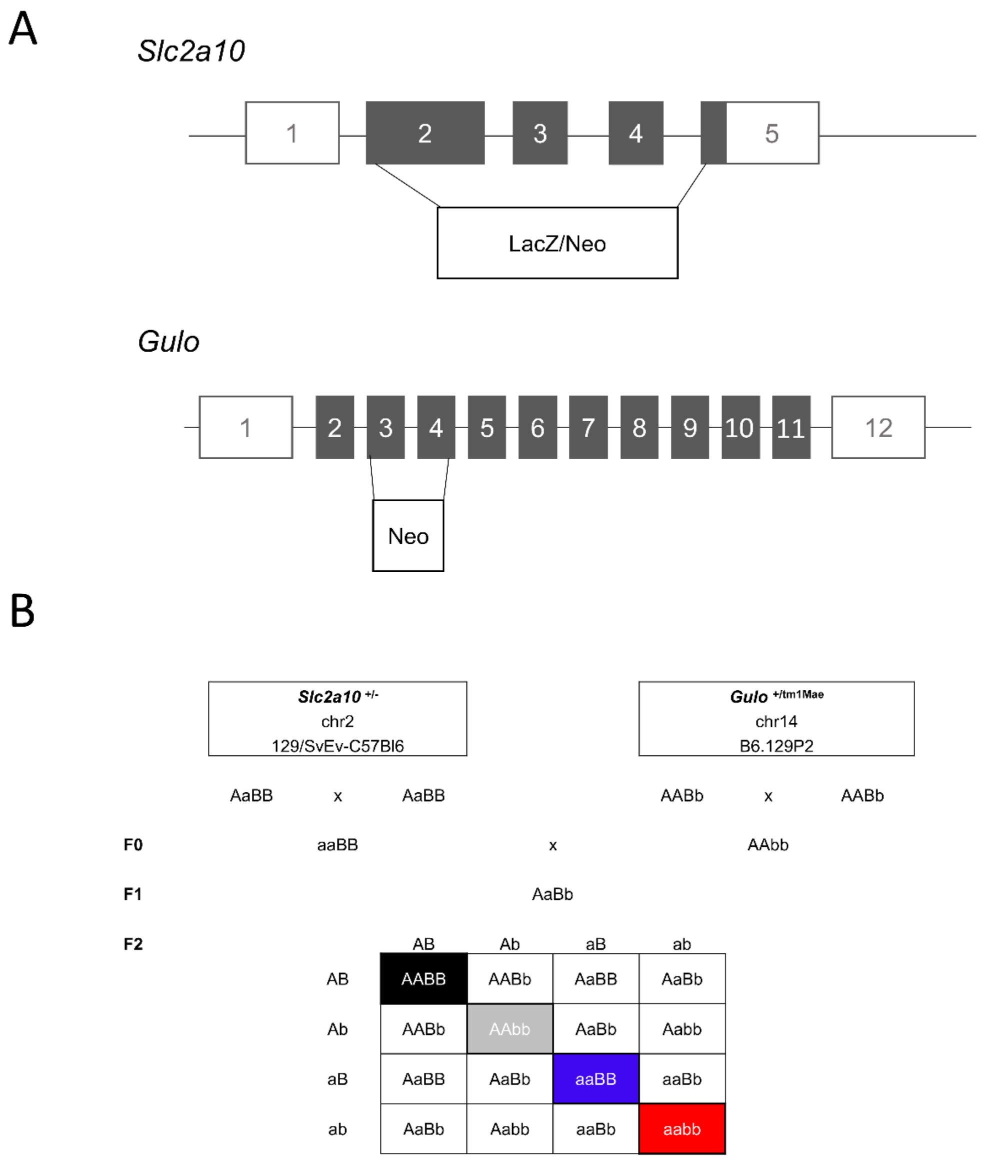


**Supplementary figure 1**: **Genetics of the novel ATS mouse model, deficient for both GLUT10 and GULO. (A)** Mutation image of the mouse models employed in this study. The *Slc2a10* constitutive knock-out model harbors a LacZ/Neo selection cassette replacing a sequence ranging from exon 2 to beginning of exon 5. The *Gulo* constitutive knock-out model harbors a Neo selection cassette replacing a sequence comprising exon 3 and 4. **(B)** Breeding scheme executed in this study. Mice, mutant for either *Slc2a10* (F0: aaBB) or *Gulo* (F0: AAbb) were crossed with each other to obtain mice heterozygous for both genes (F1). Crossing in these mice can result in pups with 9 different genotypes. Four genotypes were selected for further study: wild type (*Gulo* $^{+/+}$;*Slc2a10* $^{+/+}$ - black), single knock-out (*Gulo* $^{tm1mae/\ tm1mae}$;*Slc2a10* $^{+/+}$ - grey and *Gulo* $^{+/+}$;*Slc2a10* $^{-/-}$ - blue), double knock-out (*Gulo* $^{tm1mae/\ tm1mae}$;*Slc2a10* $^{-/-}$ - red).

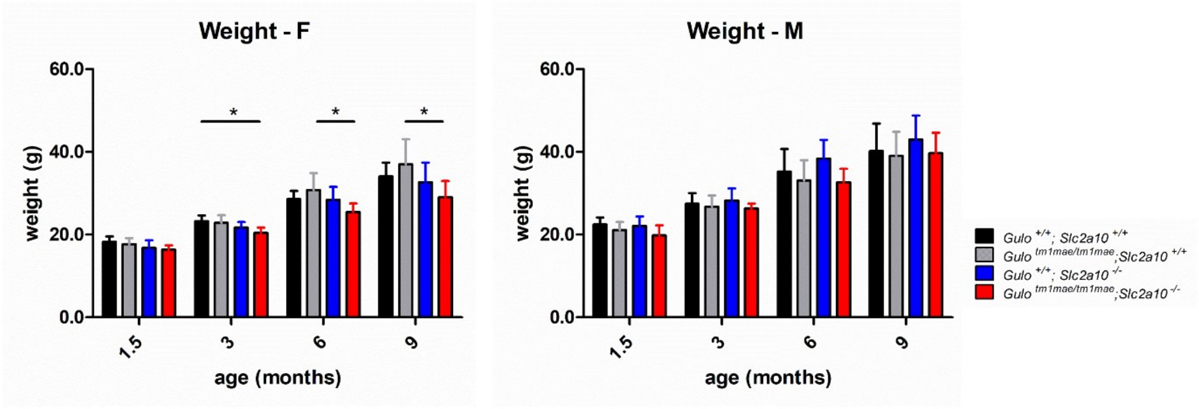


**Supplementary Figure 2: Mouse body weight.** Values plotted correspond to the means of 10 biological replicates. Error bars shown represent 95% confidence intervals. Data were analyzed using one-way ANOVA, followed by a Tukey post-hoc test. *: p ≤ 0,05, M: male, F: female

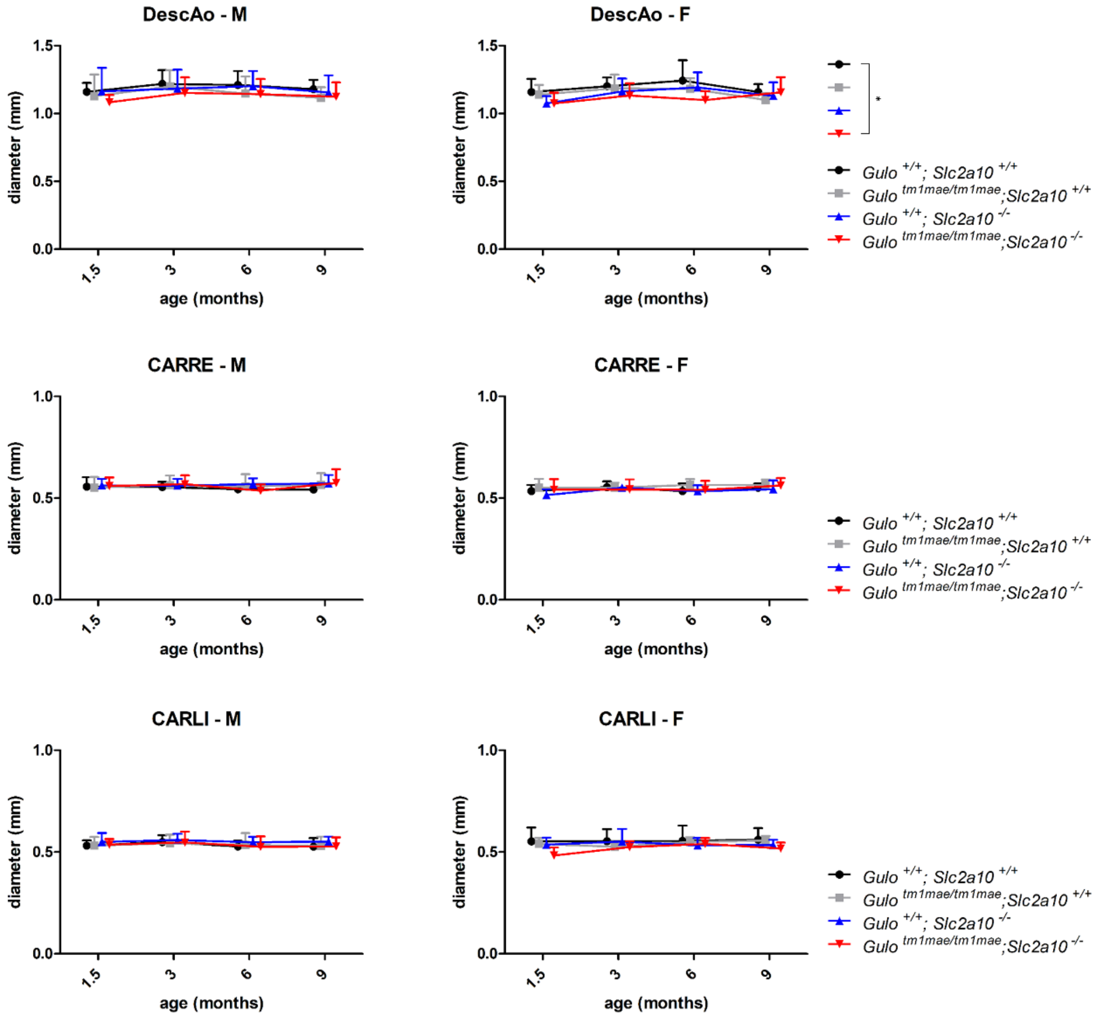


**Supplementary Figure 3: Serial ultrasound measurements obtained at 6 weeks, 3, 6 and 9 months of age at the descending aorta and carotids.** Left and right column images represent data obtained in male and female animals respectively. Shown p-values were obtained using a linear mixed model with covariance pattern modeling, without interactions. As a result, p-values should be interpreted when making the comparison for a given age and sex. Error bars shown represent 95% confidence intervals. *: p ≤ 0,05, M: male, F: female, DescAo: descending aorta, CARRE: right carotid artery, CARLI: left carotid artery

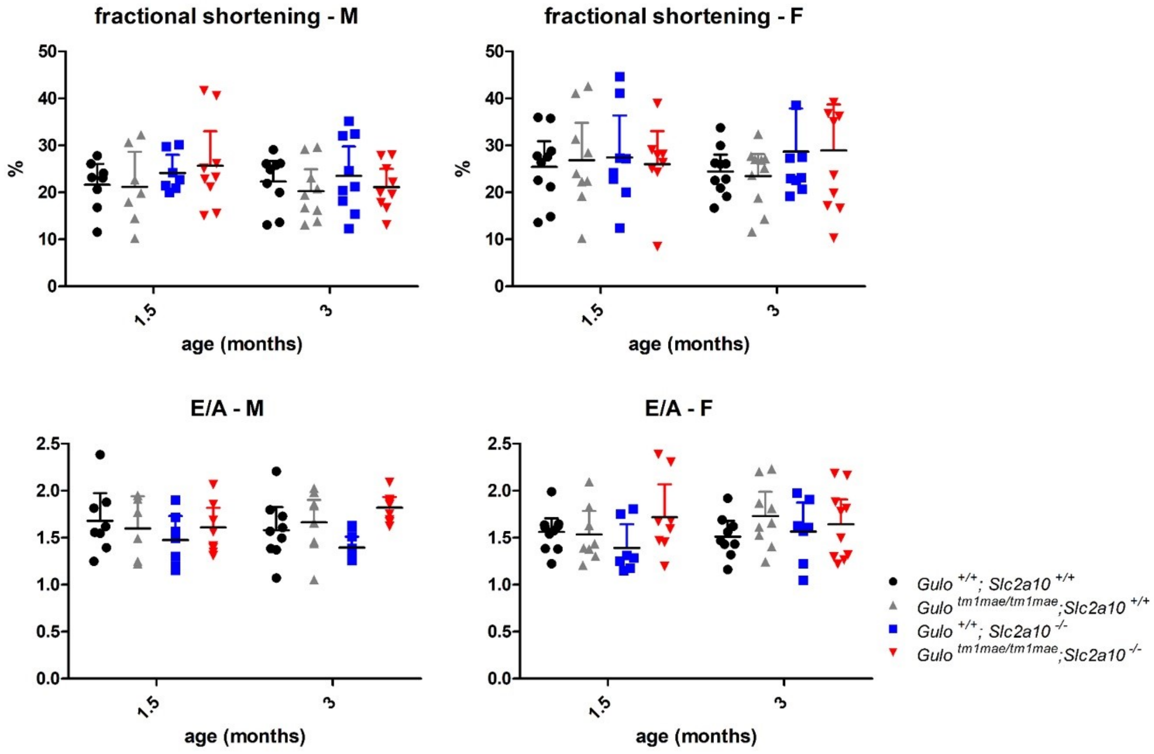


**Supplementary Figure 4: Cardiac parameters indicative for left ventricular function at 6 weeks and 3 months of age.** Error bars shown represent 95% confidence intervals. FS: fractional shortening. E/A: ratio of the early (E) to late (A) ventricular filling velocities. M: male, F: female

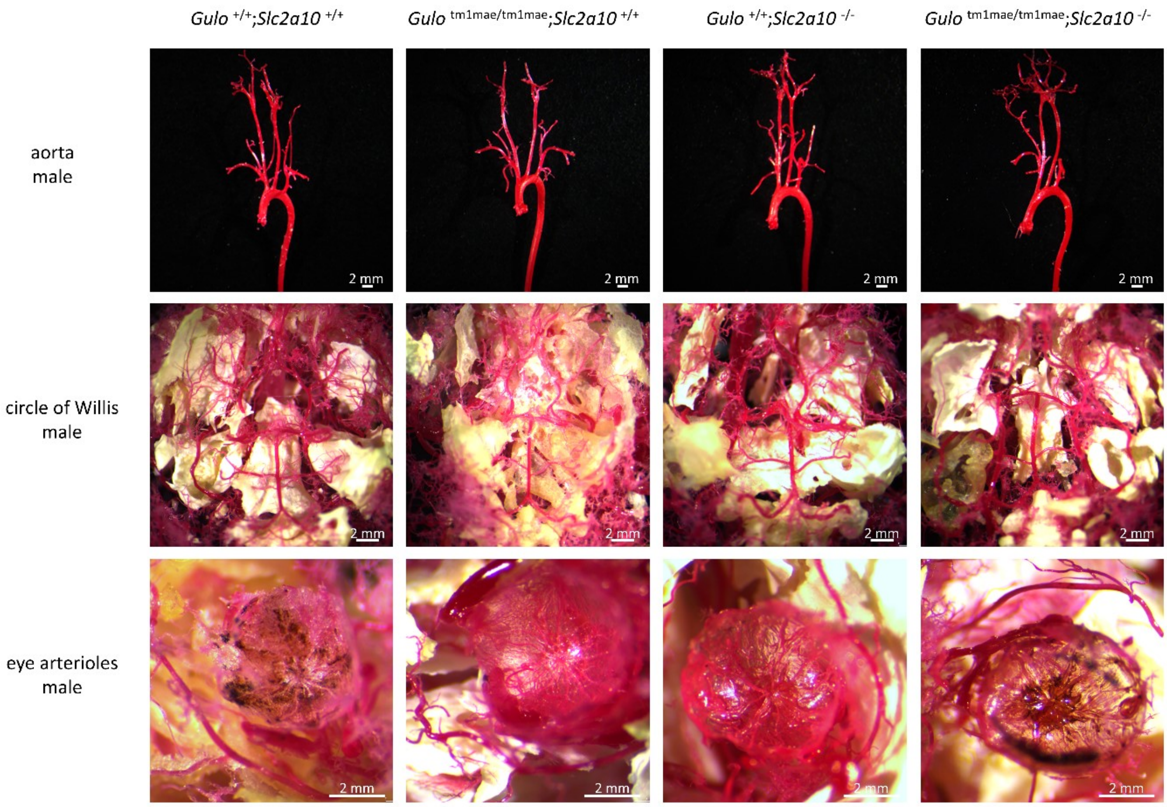


**Supplementary Figure 5: Vascular corrosion casts of *Gulo* $^{+/+}$;*Slc2a10* $^{+/+}$, *Gulo* $^{tm1mae/tm1mae}$;*Slc2a10* $^{+/+}$, *Gulo* $^{+/+}$;*Slc2a10* $^{-/-}$ and *Gulo* $^{tm1mae/\ tm1mae}$;*Slc2a10* $^{-/-}$ male mice.** Representative images of the aortic arch and its side branches, the circle of Willis and the eye arterioles are shown.

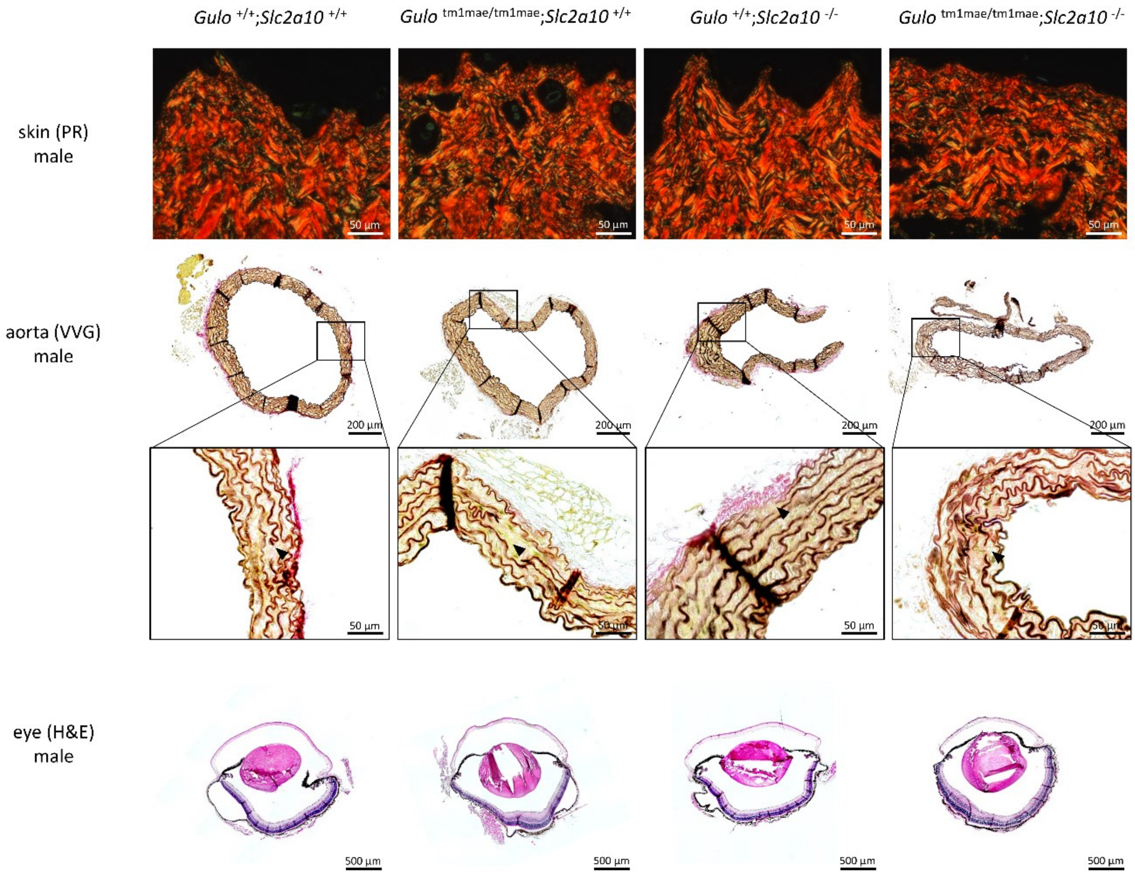


**Supplementary Figure 6: Histological analysis of *Gulo* $^{+/+}$*;Slc2a10* $^{+/+}$, *Gulo* $^{tm1mae/tm1mae}$*;Slc2a10* $^{+/+}$, *Gulo* $^{+/+}$*;Slc2a10* $^{-/-}$ and *Gulo* $^{tm1mae/\ tm1mae}$*;Slc2a10* $^{-/-}$ male mice.** Picrosirius red (PR) polarization staining (upper row) for collagen does not reveal abnormal collagen deposition in the skin of *Gulo* $^{tm1mae/\ tm1mae}$*;Slc2a10* $^{-/-}$ mice. Verhoeff-Van Gieson (VVG) elastic fiber staining (middle row) shows mild elastic fiber anomalies in the aortic wall of all studied mice (black arrows). Hematoxylin & eosin staining (bottom row) in eye tissue does not reveal major structural abnormalities in the cornea.

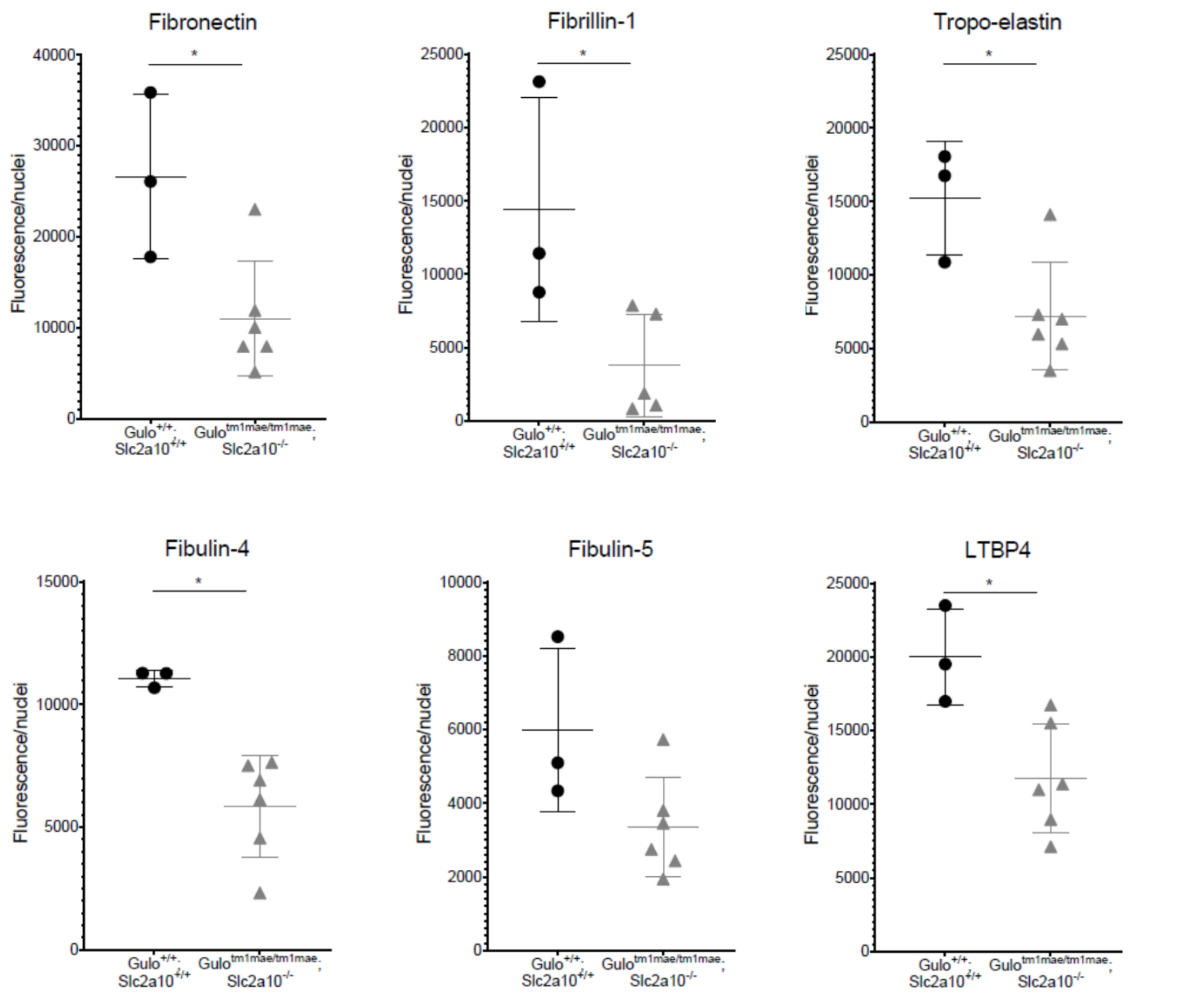


**Supplementary Figure 7: Quantification of total cell fluorescence of the ECM components corrected for the number of nuclei for the graphs, displayed in Figure 6.** Results are expressed as mean ± SD. *: $p \leq 0{,}05$.

### Supplementary tables

##### Supplementary Table 1: PCR genotyping primers and products

| *Slc2a10* | | | |
|---|---|---|---|
| **Wild type-specific product** | | **Mutation-specific product** | |
| Forward (5’-3’) | CCTTGTCGGGGGCTTCCTCATTG | Forward (5’-3’) | GCAGCGCATCGCCTTCTATC |
| Reverse (5’-3’) | CACCAGCCCCAGCCCCACTACAG | Reverse (5’-3’) | CCTCAGAGTCTCACACTCAA |
| Wild-type Band | 532 bp | Wild-type Band | none |
| Mutant band | none | Mutant band | 325 bp |
| ***Gulo*** | | | |
| **PCR with 3 primers** | | | |
| Primer 1 (5’-3’) | CGCGCCTTAATTAAGGATCC | | |
| Primer 2 (5’-3’) | GTCGTGACAGAATGTCTTGC | | |
| Primer 3 (5’-3’) | CCCAGTGACTAAGGATAAGC | | |
| Wild-type Band | 343 bp | | |
| Mutant band | 230 bp | | |

##### Supplementary Table 2: RT-qPCR primers

| *Slc2a10* | |
|---|---|
| Forward (5’-3’) | TTGGCCTGGCTTTCATCTAC |
| Reverse (5’-3’) | GCTTGTCTGAAACTGCTGCTC |
| ***ERE 1 – Rltr10b2*** | |
| Forward (5’-3’) | CCAATCCGGGTGTGAGACA |
| Reverse (5’-3’) | CTGACTCGCCAGCAAGAAC |
| ***ERE 2 – Rltr2aiap*** | |
| Forward (5’-3’) | CATGTGCCAAGGGTAGTTCTC |
| Reverse (5’-3’) | GCAAGAGAGAGAATGGCGAAAC |

##### Supplementary Table 3: Antibodies

| Primary antibody | Predicted kDa | Dilution | Manufacturer | Cat. nr. |
|---|---|---|---|---|
| Rabbit α-Fibronectin IgG | - | 1:80 | Millipore | AB2033 |
| Rabbit α-mouse Fibrillin-1 | - | 1:1,000 | Generated in Reinhardt Lab | - |
| Rabbit α-mouse Tropo-elastin | - | 1:500 | Generated in Reinhardt Lab | - |
| Rabbit α-mouse Fibulin-4 | - | 1:500 | Generated in Reinhardt Lab | - |
| Rabbit α-mouse Fibulin-5 | - | 1:500 | Generated in Reinhardt Lab | - |
| Rabbit α-mouse LTBP-4 | - | 1:500 | Generated in Reinhardt Lab | - |
| Rabbit α-SMAD2 IgG | 60 | 1:1000 | Cell signaling | 5339S |
| Rabbit α-pSMAD2 IgG | 55-60 | 1:400 | Merck Millipore | 04-953 |
| Mouse α-β catenin IgG1 | 92 | 1:2000 | BD Bioscience | 610153 |
